# Interpretation of southern hemisphere humpback whale diet via stable isotopes; implications of tissue-specific analysis

**DOI:** 10.1101/2022.02.14.480312

**Authors:** June Eggebo, Jasmin Groβ, Susan Bengtson Nash

## Abstract

Blubber and skin are commonly used tissues in stable isotope analysis for the purpose of investigating cetacean diet. Critical comparison of tissue-specific isotopic signals is, however, lacking resulting in uncertainty surrounding the representativeness and therefore utility of different tissues for accurate determination of recent foraging. This study used remotely biopsied blubber and skin tissues from southern hemisphere humpback whales for strategic comparison of δ^**13**^C and δ^**15**^N values. Samples were collected 2008-2018 as part of long-term monitoring under the Humpback Whale Sentinel Program. Blubber tissues were lipid-extracted prior to analysis, whilst mathematical lipid-correction was performed on skin samples. Isotopic values from paired blubber and skin samples from the same individuals were compared to assess whether tissues could be used interchangeably for isotope analysis and dietary interpretation. Significant differences were observed for both δ^13^C and δ^15^N, flagging previously undocumented methodological considerations, and the need for method validation and standardisation in application of these approaches. This study therefore advances methodological aspects of cetacean dietary analysis. This is of elevated importance in the context of rapidly changing ocean ecosystems.

## 1. Introduction

Southern hemisphere humpback whales (*Megaptera novaeangliae*; SHHWs) have been implemented as a sentinel species for the circumpolar surveillance of pollution and climate change in the Southern Ocean (1,2). As capital breeders, these populations rely on intensive summer feeding on Antarctic krill (*Euphausia superba;* hereafter ‘krill’; Groß et al., 2021; Waugh et al., 2012), to sustain their annual winter migrations to lower latitudes for breeding and calving. The narrow foraging niche of SHHWs, results in a distilled connection between ecosystem productivity and energetic provisioning (both prey type and foraging success; Castrillon and Bengtson Nash, 2020). Their ecophysiology thus renders these populations powerful indicators of ecosystem productivity and change.

Krill are a sympagic species, with sea-ice providing feeding habitat and refuge for early life stages (6,7). Polar ecosystems are undergoing rapid change, manifesting in sea-ice melt (8), ocean acidification (9), and a rise in sea water temperature (10). These physio-chemical characteristics of the krill ecosystem, in turn, impact krill recruitment and survival (11,12). Any change in the abundance or availability of krill is expected to carry significant implications for krill consumers (Tulloch et al 2019, Seyboth et al. 2016). Humpback whale (HW) populations globally show a high degree of plasticity in both their target prey and foraging behaviour (13,14). As such, SHHWs may be expected to respond to a change in krill availability through diversified foraging, including changes to both prey and foraging range. Longitudinal monitoring of SHHW diet has therefore been identified as a core sentinel parameter under the Humpback Whale Sentinel Program (HWSP), with interannual variation and drift assumed to reflect a change in krill availability.

Ecologists use bulk stable isotope analysis (BSIA) to directly identify and trace elemental cycling in the biosphere (Fry, 2008). Over the last few decades, BSIA has played a significant role in research involving animal migration (15,16), diet (17,18), reproduction (19,20) and food web connectivity (21,22). The stable isotopes of carbon (^13^C,^12^C) and nitrogen (^15^N,^14^N) have, in particular, become valuable in diet research of marine mammals (23–26). The use of stable isotope analysis to investigate the structure of food webs is based on two assumptions: namely that the isotopic composition of consumer tissue reflects the isotopic composition of their diet, and that consumers are slightly enriched in ^15^N and to a lesser extent in ^13^C compared to their food (27,28). The phenomenon is called ‘trophic discrimination’, also referred to as ‘trophic fractionation’ and averages 0.5-1.0 ‰ for carbon (Δ^13^C; DeNiro and Epstein, 1978; Fry, 2008; Zuev et al., 2019) and 2-4 ‰ for nitrogen (Δ^15^N; Minagawa and Wada, 1984; Fry, 2008; Zuev et al., 2019). Trophic levels (TLs) are a hierarchical way of classifying organisms according to their theoretical feeding relationships within an ecosystem (32). Nitrogen isotopes (δ^15^N) increase as a function of mean TL (Minagawa and Wada, 1984) due to the relatively faster metabolic loss of ^14^N compared to ^15^N leaving animals at higher trophic levels with higher δ^15^N values (Fry, 2008). Carbon isotopes (δ^13^C) in marine environments can be traced from producers such as particulate organic matter (POM) and phytoplankton, to consumers to determine primary carbon sources (27,33). These values are often used to distinguish between two geographically distinct food webs. Altabet and Francois (1994) demonstrated that surface water δ^13^C values of POM lay at approximately −22 ‰ in temperate latitudes but decrease to −25 ‰, sometimes down to −35 ‰ (35) closer to Antarctica. Thus, animals feeding in Antarctic food webs demonstrate correspondingly low carbon isotope values (36–39), compared to those feeding temperate food webs (40,41).

The interpretation of bulk stable isotope (BSI) signals is, however, not without uncertainty. In addition to the prey type and foraging range, the trophic position (TP) is known to be influenced by endogenous factors such as nutritional stress, metabolic activity of tissues, diet quality, body size, excretory mechanisms and feeding rate (42–46). Further, the isotopic signals associated with tissues of different biomolecular composition (i.e. lipids, carbohydrates, proteins) have frequently been recorded (30,46,47). The extent to which tissue types within an individual differ in their δ^15^N and δ^13^C values thus carries inherent uncertainty for robust quantification of diet and represents a methodological aspect of cetacean dietary investigation that has not been thoroughly addressed.

In cetacean research, blubber and skin tissue are the most commonly used tissue types for dietary investigation as they are metabolically active and can easily be obtained via non-lethal biopsies from healthy, free-swimming individuals (48,49). Marine mammal blubber is principally composed of lipids and contains small amounts of protein (46,50). By contrast, skin mainly contains protein and a limited amount of lipids (46,51,52). In BSIA, lipids confound analyses by decreasing the tissue ^13^C/^12^C and hence lowering measured δ^13^C values (30). As such, the influence of lipid content on whole tissue δ^13^C values, and following dietary interpretation must be accounted for (53). Two approaches are commonly used to account for lipids. The first methods is the physical removal of lipid fractions through solvent extraction prior to BSIA. Alternatively, where the relationship between lipid-containing and lipid-depleted tissues of a species is known, mathematical corrections have been developed and applied (Post et al., 2007; Ryan et al., 2012, Groß et al. 2021).

In an effort to further strengthen data obtained from long-term monitoring of SHHW diet, the current study sought to compare the BSI measurements obtained from lipid-adjusted blubber and skin tissues respectively. In order to test the hypothesis that δ^13^C and δ^15^N values of blubber and skin taken from the same individual could be used interchangeably, 171 paired samples were investigated, providing new insights into method application, data interpretations, and species physiology.

## 2. Material and methods

### 2.1 Sample Collection

Blubber and skin biopsy samples were obtained for long-term monitoring under the HWSP from free-swimming SHHW of the east coast of Australia-migrating stock (E1 as defined by the International Whaling Commission; Acevedo et al., 2013), between 2008 and 2018. The biopsies were collected off North Stradbroke Island, southeast Queensland, Australia (approximately 27°26 S, 153°34 E) during the annual northward (June/ July) and southward (September/ October) migration according to methods described by Bengtson Nash et al. (2018). The collection of samples was carried out under Scientific Purposes permit, granted by the QLD department of Environment and Heritage Protection and animal ethics permit granted by the Griffith University Animal Ethics Committee. In total, 171 paired blubber and skin biopsy samples were included in this study. Blubber tissue was lipid extracted with solvents prior to analysis while skin tissue was mathematically lipid corrected. Both are referred to as “lipid-adjusted” in subsequent text.

### 2.2 Lipid Adjustment

#### 2.2.1 Solvent Extraction

Approximately 30 mg of blubber was lipid extracted prior to BSIA. The solvent lipid extraction of blubber tissue was completed using a modified methanol-dichloromethane-water (2:1:0.8 v/v/v MeOH/CH_2_Cl_2_/H_2_O) method pioneered by Bligh and Dyer (1959), as described in detail elsewhere (e.g. Groß et al., 2021).

#### 2.2.2 Mathematical Correction

Previously, Groß et al. (2021) determined the most appropriate isotopic discrimination factor of skin for the study population to be 8.92 ‰. The mass balance approach (MBA) developed by Fry (2002), was considered the best fit for lipid correction of SHHW skin, and was therefore applied in the current study. The correction applied in this study was as follows (Eq. 1):

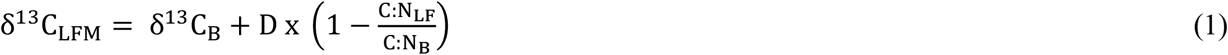

Where δ^13^C_LFM_ is the lipid-free (or lipid-corrected) carbon isotope value of skin, δ^13^C_B_ is the bulk carbon isotope value measured from Balaenopteridae skin, and D is the isotopic discrimination factor. C: N_LF_ is the measured ratio of lipid-free skin tissue, whilst C: N_B_ is the measured ratio of bulk Balaenoptera skin tissue.

### 2.3 Bulk Stable Isotope Analysis

Lipid-adjusted blubber and skin tissue were oven dried overnight at approximately 58°C and pulverized in to 1-2 mg samples which were placed into tin capsules for δ^13^C and δ^15^N analysis. Stable isotope abundances were calculated in permil using the following equation:

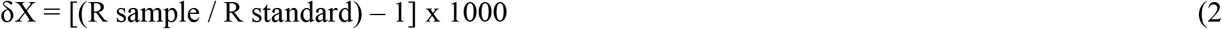

Where, X is ^13^C or ^15^N, and R is the respective ratio ^13^C/^12^C or ^15^N/^14^N. The international reference standards used for carbon and nitrogen are, respectively, Vienna Pee Dee Belemnite and N_2_ in air. Laboratory standards, sucrose and (NH_4_)_2_SO_4_ were calibrated using international standards IAEA-CH_6_ for carbon and IAEA N1 for nitrogen. The preparation system used is a Europa EA-GSL interfaced to a SERCON Hydra 20–20 isotope ratio mass-spectrometer (IRMS). Based on analysis of replicate standards, the standard deviations for δ^13^C and δ^15^N averaged 0.1 ‰ and 0.15 ‰, respectively.

### 2.4 Krill range calculation

The krill range i.e., the isotopic range expected for individual whale δ^13^C and δ^15^N values feeding exclusively on Antarctic krill, was calculated based on isotopic values of krill derived from Eisenmann et al. 2016. Blubber and skin trophic fractionation (TF) estimates were calculated in this study (Table S1, Table S2). The krill range for lipid-extracted blubber was −28.14 to −24.66 and 5.96 to 9.34, while for lipid-corrected skin the range was −27.09 to −23.61 and 5.12 to 8.50 respectively for δ^13^C and δ^15^N. This facilitated comparison of blubber and skin foraging results, allowing for an inter-annual evaluation of diet representation within and between tissues throughout sample years.

### 2.5 Trophic position calculation

Trophic position for SHHWs was calculated from lipid-adjusted blubber and skin tissues, relative to krill using the following equation:

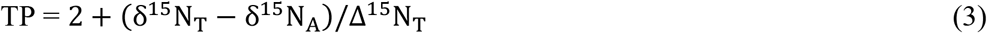

Where, 2 is the TP of the primary consumer, T is tissue type (lipid-extracted blubber or lipid-corrected skin), A is Antarctic krill (prey) and Δ^15^N is the TF value. Mean lipid-corrected skin and lipid-extracted blubber δ^15^N isotope values were derived from BSIA in this study, the mean Antarctic krill δ^15^N value of 3.2 ‰ was derived from literature estimates as shown in Table S1.1, and TF values for lipid-extracted blubber (4.45 ‰) and lipid-corrected skin (3.61 ‰) were calculated as shown in Table S1.2.

### 2.6 Statistics

Data analyses were performed in R version 1.3. 1093 (61) and GraphPad Prism version 9.0.2 (62). A Shapiro-Wilk test and a Levene’s test were used to test the data for normality and homogeneity of variance, respectively. All statistical results were interpreted using a significance level of α = 0.05. The δ^13^C and δ^15^N isotopic values across sex and migration showed no significant difference (p=0.1841 and p=0.1184 respectively), thus all samples were treated as a homogenous cohort. A Shapiro-wilks test demonstrated non-normality for δ^13^C and δ^15^N isotopic values within and between lipid-adjusted blubber and skin, thus non-parametric statistical tests were further applied. Two separate Wilcoxon matched pair signed rank tests were used to test for differences in δ^13^C and δ^15^N values between the two tissue types. The test structure used δ^13^C and δ^15^N as test variables for differences in the factor ‘tissue type’ with fixed values for lipid-adjusted blubber and skin tissue. A non-parametric Kruskal-Wallis test with multiple comparisons was applied to investigate trends across sample years for δ^13^C and δ^15^N values.

## 3.0 Results and Discussion

The present study is the first to investigate tissue specific BSI measurements and implications for interpretation of SHHW diet. Our results showed that there are significant differences in δ^13^C and δ^15^N values obtained from lipid-adjusted blubber and skin from the same individuals. Such differences were more prominent in some individuals, thus occasionally led to different down-stream interpretation of trophic position. There was greater variability in δ^15^N values of lipid-extracted blubber compared to lipid-corrected skin. The tissue-specific variation in δ^15^N values was surprisingly not reflected in tissue-specific TP estimates as lipid-adjusted blubber and skin tissue demonstrated a similar TP of 3.0. These findings underscore that tissue-specific variation must be thoroughly investigated before comparing dietary results obtained via BSIA using two different tissues and caution against interchangeable use of tissues or comparison between them.

### 3.1 Bulk differences

For both δ^13^C and δ^15^N values of lipid-adjusted tissues, significant differences were observed (δ^13^C p=0.0001 and δ^15^N p=0.0001; Figure 1). Lipid-extracted blubber values showed greater variability for both δ^13^C and δ^15^N compared to lipid-corrected skin (Table 1., Figure 1).

**Table 1:**
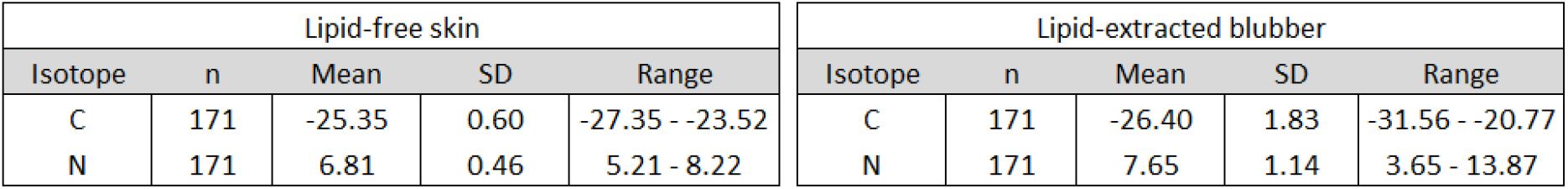
Table overview of the mean, standard deviation (SD) and range for δ13C and δ15N values for lipid-extracted blubber and lipid-corrected skin tissue of E1 humpback whales (n=171).

| Lipid-free skin |  |  |  |  | Lipid-extracted blubber |  |  |  |  |
| --- | --- | --- | --- | --- | --- | --- | --- | --- | --- |
| Isotope | n | Mean | SD | Range | Isotope | n | Mean | SD | Range |
| C | 171 | -25.35 | 0.60 | -27.35 - -23.52 | C | 171 | -26.40 | 1.83 | -31.56 - -20.77 |
| N | 171 | 6.81 | 0.46 | 5.21 - 8.22 | N | 171 | 7.65 | 1.14 | 3.65 - 13.87 |

**Figure 1:**
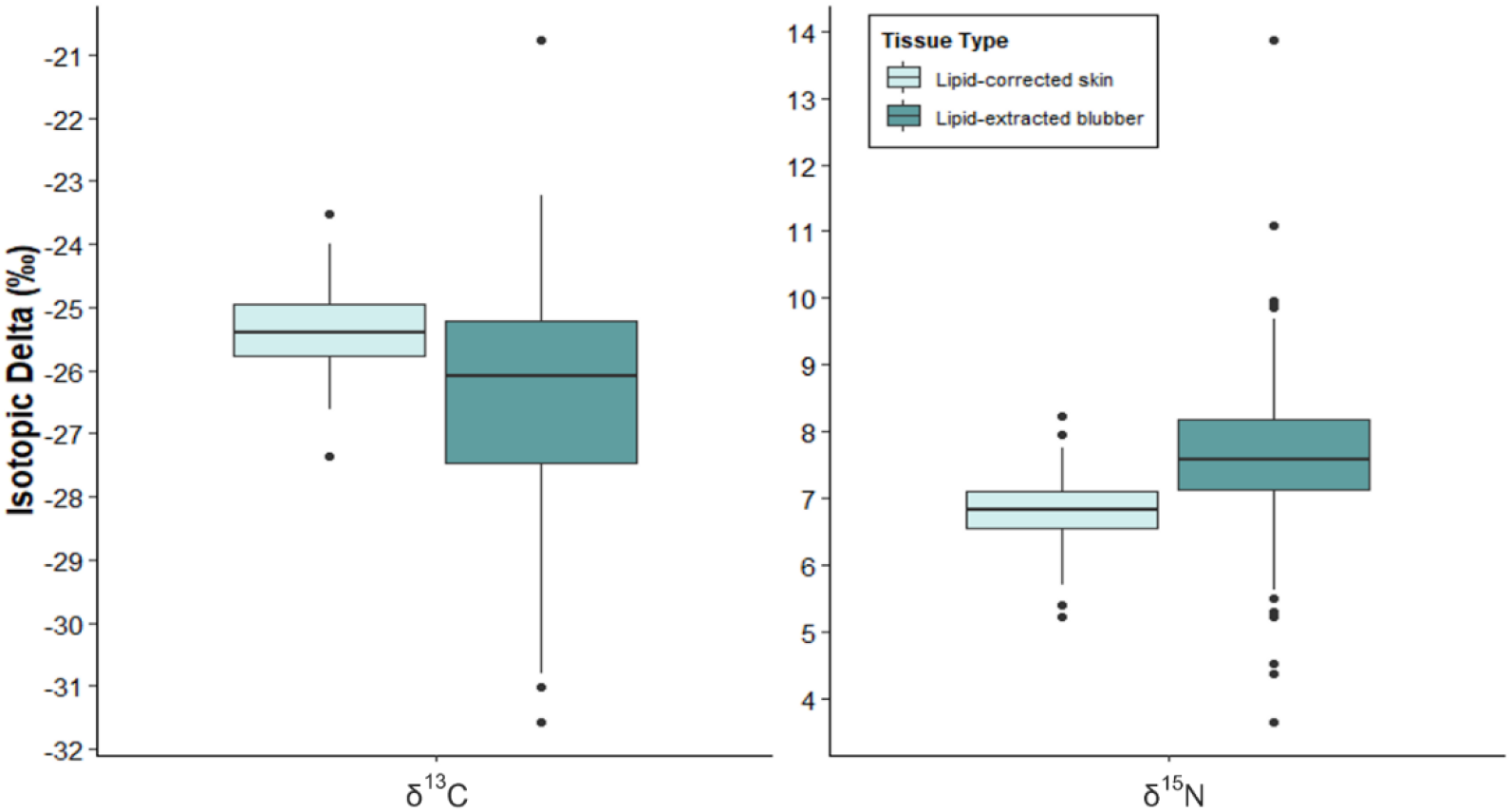
Box plot showing the distribution of δ13C and δ15N values for lipid-extracted blubber and lipid-corrected skin tissue (n=171).

As the tissues were obtained from the same individual whale, the extent of the variability in both isotope signatures was not expected. There is limited research on the comparison of δ^13^C and δ^15^N values between HW blubber and skin tissue, however, a significant difference between the two tissues either for one or both isotopes has been documented (e.g Groß et al., 2021; Todd et al., 1997). However, the reasons for this variation are not clear, and thus we attempt to evaluate several factors that may have contributed to the significant differences found in this study.

### 3.2 Inter-annual differences

Large inter-annual variability in isotopic signatures has previously been evidenced via fatty acid analysis for this population (Groß et al. 2020). When samples were separated by year, limiting analysis to those years where >10 paired samples were available for analysis (2013-2018), significant inter-annual differences were observed in selected years. Of the six within year cohorts available, three demonstrated a significant difference in the δ^13^C values between the two tissue types (2016; p=0.0216, 2017; p=0.0335 and 2018; p=0.0001; Figure 2A). Similarly, three years, albeit three different years, showed significant differences in δ^15^N values between tissue types (2013; p=0.0003, 2014; p=0.0001 and 2017; p=0.0001; Figure 2B).

**Figure 2:**
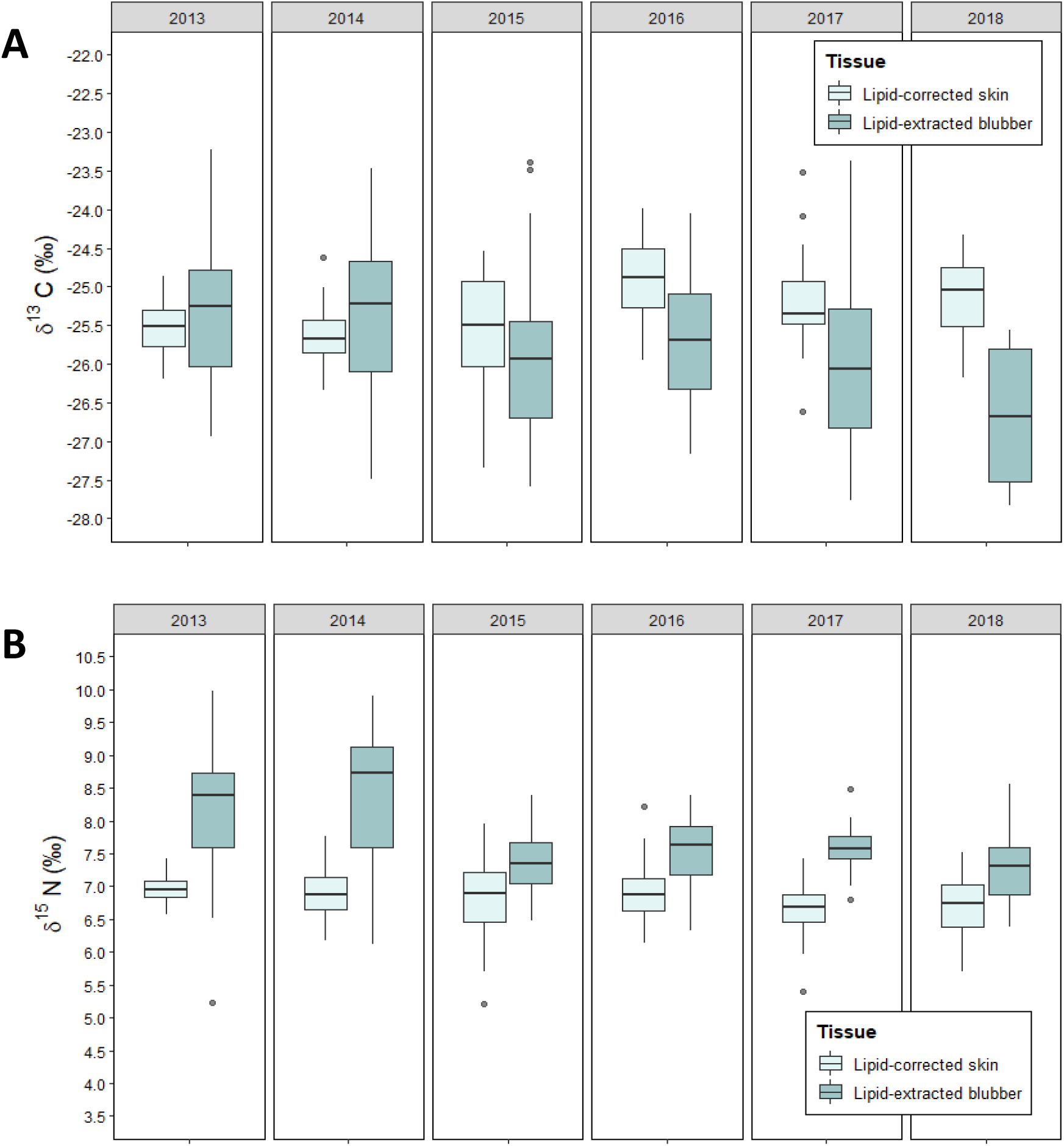
Isotopic values of blubber and skin (n=171) across all sample years. (A) illustrates comparison between both tissues for δ13C and (B) for δ15N values.

As the significant differences in δ^13^C and δ^15^N values between lipid-adjusted blubber and skin did not occur in the same sample years, there may be underlaying tissue-specific variations that could be driving the variability in isotope signatures. Figure 2 illustrates an overall low variability in both δ^13^C and δ^15^N values for lipid-corrected skin across all sample years, compared to lipid-extracted blubber that has a greater variability with more prominent oscillations in some years like 2014. The differences in isotopic signatures between the tissues may lead to issues for interpretation because we cannot be certain whether the variability present in blubber δ^13^C and δ^15^N values is caused by variability in prey type or foraging location, or whether the observed variability is introduced by endogenous factors or method artefacts. Hence, we do not know if we lose information about foraging variability when we just interpret results from skin, or if we introduce variability to results when we just interpret results from blubber tissue.

### 3.3 Trophic position comparison

Trophic position estimates were calculated to investigate whether the observed differences between blubber and skin δ^13^C and δ^15^N values also leads to differences in the dietary information derived from the two tissue types. Overall, when samples were considered as a single cohort, TP did not vary significantly between the two tissues (Wilcoxon p=0.988). The mean TP was 3 ± 0.26 for blubber and 3 ± 0.13 for skin. However, as with single year cohorts for BSI values, significant differences in tissue-specific TP were found in the years 2013 (p=0.0211), 2014 (p=0.0113) and 2016 (p=0.0218; Figure 3).

**Figure 3:**
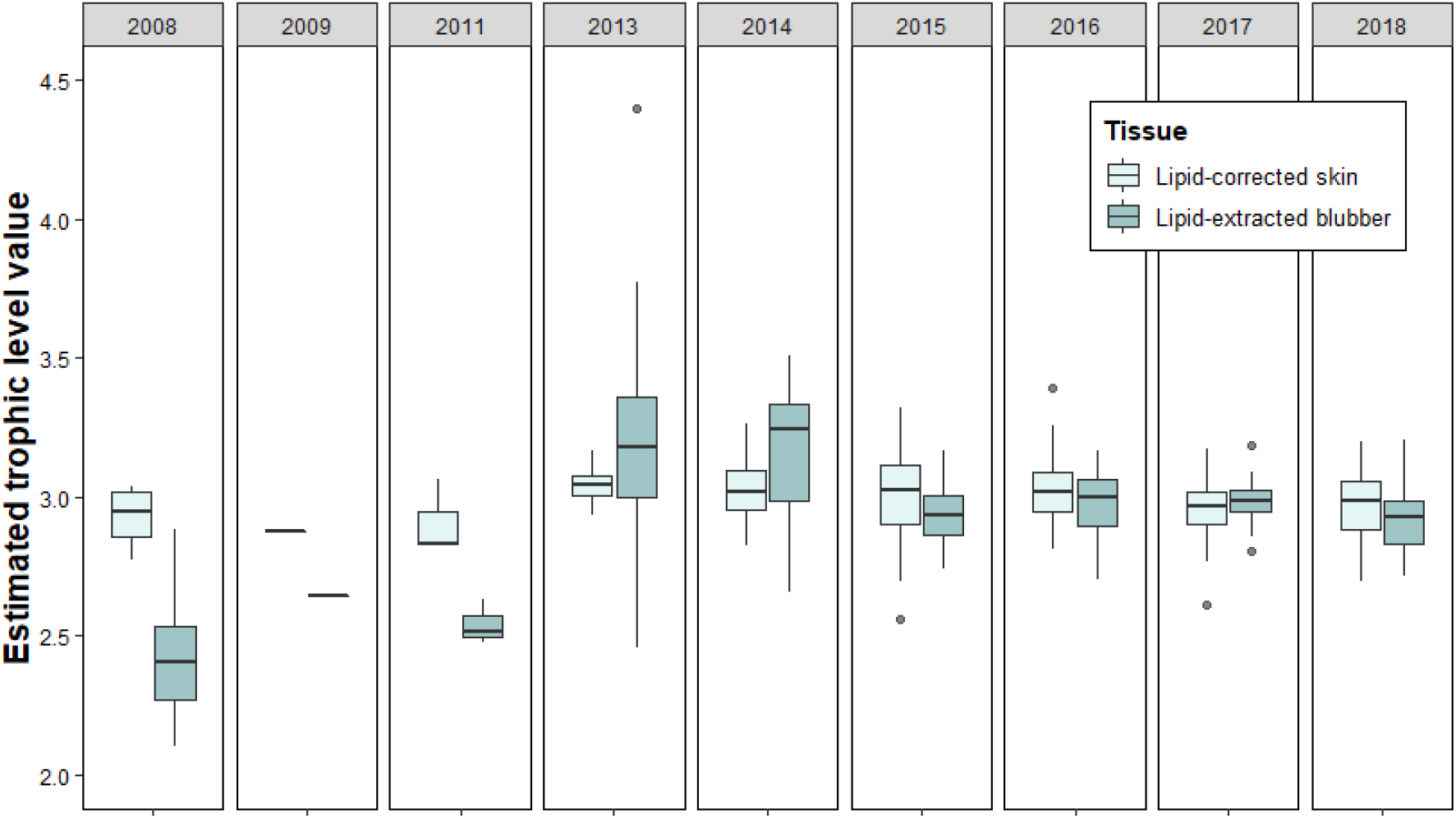
Trophic position estimates for blubber and skin tissue across all sampling years.

Although there was an overall significant difference in δ^15^N values between lipid-adjusted blubber and skin tissue, this difference was not reflected in the TP values calculated for both tissues (TP = 3.0). The calculated TP values are congruent with the classical feeding paradigm of a high-fidelity krill diet in SHHWs (63–66). However, as some variation was observed in 2013, 2014 and 2016 (Figure 3), there are small, but minor underlying differences that reflect the significant differences in δ^15^N values between the two tissues.

The equation used to estimate TP has limitations, which can lead to errors in interpretation. First, the trophic fractionation factors (Δ^15^N) used in the equation, 4.45 ‰ for lipid-extracted blubber and 3.61 ‰ for lipid corrected-skin tissue, are only based on estimates. The true TP for E1 humpback whale tissues are unknown. Additionally, Δ^15^N vary between and within species and tissues, introducing error when estimates are based on other tissues or species. Secondly, an average δ^15^N value for krill was used in the equation, which introduces errors as there are spatial and temporal differences in δ^15^N values of krill (67–70). Some introduced uncertainty could be reduced by analysing compound specific nitrogen isotope composition of amino acids (68–70), however we were unable to analyse compound specific isotopes results due to cost restrains.

### 3.4 Tissue-specific krill space

The implications of tissue-specific variability in BSI values for the interpretation of diet was further investigated by creating a krill space (isotope range) for each tissue. The shaded areas in Figure 4 illustrate the tissue-specific krill space in which SHHW δ^13^C and δ^15^N values are expected to fall if the individual whales were feeding primarily on krill the austral summer prior to sampling. The figure only shows the δ^13^C and δ^15^N values of lipid-adjusted blubber and skin from two sample years, 2013 and 2015, as these years highlight the two different scenarios that we have observed between 2008 and 2020; lipid-corrected skin isotope values fall within the calculated krill space while either the majority of both δ^13^C and δ^15^N values of lipid-extracted blubber fall outside the krill space or the majority of just δ^13^C values of blubber fall outside the krill space.

**Figure 4:**
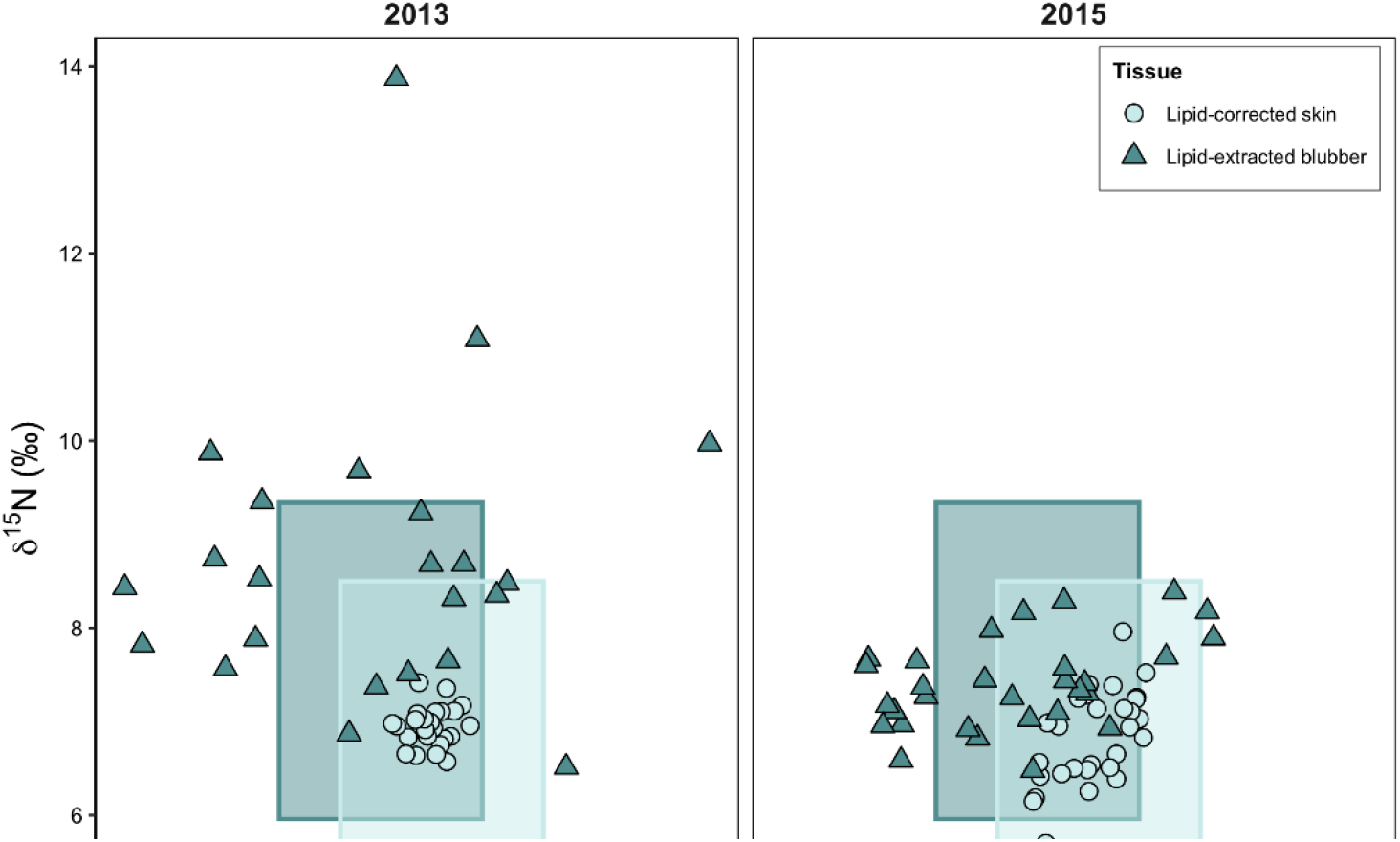
Scatterplot illustrating the tissue-specific krill space for δ13C and δ15N values of lipid-adjusted blubber and skin tissue of 2013 (n=24) and 2015 (n=30).

For lipid-extracted blubber, in 2013 only 29.2% and 53.3% of the isotopic data points fell within the krill space in 2013 and 2015, respectively. This was half of what was observed for lipid-corrected skin, where 100% of the data fell within the krill space in 2013, and 96.7% in 2015. This leads to different interpretations about the diet plasticity of SHHW. If we would make inferences based on skin isotope results, we would conclude that SHHW exclusively feed on krill in the Southern Ocean. However, if we would only interpret blubber isotope results, we would conclude that SHHW exhibit a much greater diet plasticity than expected for a high-fidelity krill diet species. Interestingly, the observed variability does not translate into different interpretations of TP.

### 3.5 Factors influencing variability

#### 3.5.1 Endogenous factors

To properly interpret stable isotope signatures of animal tissues, it is essential to account for temporal dynamics of isotopic integration such as tissue turn-over rate and diet-tissue incorporation.

Isotopic turnover time describes the time it takes for a tissue layer to be replaced entirely by a new layer of tissue (71,72). It is an important consideration when assessing different tissues as isotopic incorporation occurs during tissue growth, resynthesis and breakdown, and can vary among tissue types (47,73). The turnover time for blubber and skin of SHHW is unknown, however SHHW blubber turnover is suggested to be <9 months, because the blubber lipid store is almost entirely depleted over the course of their annual migration, due to prolonged fasting (66). Isotopic turnover time for skin δ^15^N has been estimated to be approximately 180 days for bottlenose dolphins (*Tursiops truncates*) and 163 days for blue whales (*Balaenoptera musculus*), while δ^13^C has been estimated to be approximately 104 days for bottlenose dolphins (74,75). Based on taxonomy and size, we therefore expect SHHW skin to have an isotopic turnover time that ranges from approximately 104 to 180 days. In the present study, there were roughly 60 days between E1 humpback whales leaving their Antarctic feeding grounds in March and the time they were sampled in June/ July, and roughly 150 days until they were sampled in September/ October. Although, turnover time for either SHHW tissue is unknown, we can assume that both tissues reflect a similar diet intake timeframe based on available information.

Aside from tissue turnover differences, the variation in δ^13^C and δ^15^N values between lipid-adjusted skin and blubber tissues may be linked to differences in tissue-specific metabolic routing, which is expected to produce a consistent offset between the stable isotope values of individual tissues. Metabolic routing of different biomolecules during tissue synthesis and metabolism impacts diet-tissue isotope discrimination. This means that some tissues may primarily reflect individual diet components such as carbohydrates and lipids derived from one dietary source and proteins derived from another (76,77). By way of example, a study by Misra et al. (2019) on bottlenose dolphins found that blubber tissue likely represents metabolic patterns linked to fatty acids and ketogenic amino acids related to fat synthesis and deposition within the tissue, whilst skin showed metabolites involved in gluconeogenic pathways pointing to active anabolic energy-generating metabolism. By extension, it is possible that the δ^13^C and δ^15^N values of SHHW blubber tissue may be drawn from a more direct energy pool, where lipids are immediately stored in the blubber, while secondary pathways may be involved in the growth of skin tissue. The complexity of tissue-specific metabolic routing and discrimination can also lead to uncertainty in lipid normalization models due to unknown protein-lipid discrimination values.

#### 3.5.2 Artefacts

In addition to endogenous factors, methodological artefacts should also be considered as a source of variation. The observed differences between both tissues may be related to the lipid-adjustment approaches applied to the respective tissue type. The mass-balance mathematical lipid correction model proposed by Fry (2002) relies on precision, accuracy and reliability in predicting the lipid-free δ^13^C values. The model is based on C:N ratios and thus lipid content, which was estimated to have a mean standard error of ~0.05 in predicting lipid-free δ^13^C values for skin tissue of E1 humpback whales (3). A study by Groß et al. (2021) specifically calculated the discrimination value ‘D’ for skin tissue of individual E1 humpback whales to be applied in the mass balance correction model, which gave a ‘D’ value of 8.92 ‰ and a C:N_LM_ of 3.1. The authors recommended the use of these values in conjunction with the mass balance model for E1 humpback whale skin tissue, if the skin tissue has a low lipid content, leading to small lipid corrections that limit errors in interpretation. However, the use of these exact coefficient values for ‘D’ and C:N_LM_ increases uncertainty if the correction is applied to species and populations where empirical values are unknown. Thus, although the ‘D’ value has been determined for E1 humpback whales, the accuracy of the value is unknown, given the large interannual variability in δ^13^C and δ^15^N values (e.g. Bengtson Nash et al., 2017; McConnaughey and McRoy, 1979; Tieszen et al., 1983).

As with mathematical lipid correction, solvent-extraction may be similarly susceptible to the introduction of methodological artefacts. The dichloromethane/methanol solvent combination used in this study has been reported to have little influence on δ^15^N values (80). However, previous studies using various solvent combinations for lipid extraction have detected fluctuations in δ^15^N values as a result of solvents interfering with structural components of the tissue (54,81,82). This could be linked to the high δ^15^N values found in this study where 23% of all lipid-extracted blubber δ^15^N data falls outside the 95% confidence interval. In addition, all figures indicate a high variability and range in lipid-extracted blubber tissue. An increase in δ^15^N resulting from solvent lipid extractions has been linked to the loss of nitrogenous components such as amino acids (AA), which may be extracted unintentionally from the tissue as the solvents can remove polar and non-polar compounds in the process (83). The hypothesis is that methanol, which removes mostly polar structural fat components that are attached to proteins, also removes amino acids at the same time as structural fats, resulting in enrichment of ^15^N (Sotiropoulos et al., 2004 and references therein). Although this study did not seek to address this method component, altered δ^15^N values post extraction have previously been observed in fish tissues; muscle and whole body samples (84), and liver tissues (80). A study by Ryan et al. (2012) found significant increases in δ^15^N values post lipid extraction for blubber of fin whales and skin of minke whales (*Balaenoptera acutorostrata*), where the overall changes were more prominent in blubber than skin tissue, which is logical given the respective lipid proportions. Thus, we hypothesis that E1 humpback whale blubber, being an adipose tissue with high lipid content is susceptible to solvent extraction related removal of amino acids resulting in the possibility of distorting the signal of δ^15^N values in BSIA.

## 4. Conclusion

This study showed that the overall comparison of lipid adjusted blubber and skin δ^13^C and δ^15^N values of SHHW were similar, but not to the extent that we can confidently recommend the interchangeable use of both tissues in this field of research. Although the mean trophic position of each year cohort was similar, the greater variability observed in blubber, which may be interpreted as higher trophic level feeding, is not present in skin values. This variability has been related to variation in lipid content, solvent interference, isotopic discrimination, and metabolic pathways between blubber and skin tissue. All are key factors that can impact the interpretation of stable isotope results. We recommend that future studies incorporate a standard for SHHW blubber and skin tissue, with the application of multiple lipid standardization approaches. Additionally, we suggest the inclusion of multiple solvent lipid extraction trials for blubber tissue to determine the potential impact on isotopic signatures. This will allow for optimization of dietary investigation and standardization of methodologies, which will improve long-term monitoring of SHHWs to provide new insights into energy utilisation by these populations.

## Acknowledgements

This work was funded by a Griffith University Honours Student Grant. June Eggebo thanks the Centre for Planetary Health and Food Security for her Honours Thesis Write-up Scholarship. The authors acknowledge the support from the Griffith University Stable Isotope Laboratory staff and the contributions of field volunteers who assisted with sample collection.

## References

1. Bengtson Nash SM, Castrillon J, Eisenmann P, Fry B, Shuker JD, Cropp RA, et al. Signals from the south; humpback whales carry messages of Antarctic sea-ice ecosystem variability. Glob Chang Biol [Internet]. 2017 Apr 1 [cited 2020 Jul 22];24(4):1500–10. Available from: http://doi.wiley.com/10.1111/gcb.14035

2. Druskat A, Ghosh R, Castrillon J, Bengtson Nash SM. Sex ratios of migrating southern hemisphere humpback whales: A new sentinel parameter of ecosystem health. 2019 [cited 2020 Jul 27]; Available from: https://doi.org/10.1016/j.marenvres.2019.104749

3. Groß J, Fry B, Burford MA, Bengtson Nash S. Assessing the effects of lipid extraction and lipid correction on stable isotope values (δ^13^C and δ^15^N) of blubber and skin from southern hemisphere humpback whales. Rapid Commun Mass Spectrom. 2021;35(16): 1–11.

4. Waugh CA, Nichols PD, Noad MC, Nash SB. Lipid and fatty acid profiles of migrating Southern Hemisphere humpback whales *Megaptera novaeangliae*. Mar Ecol Prog Ser. 2012;471:271–81.

5. Castrillon J, Bengtson Nash S. Evaluating cetacean body condition; a review of traditional approaches and new developments [Internet]. Vol. 10, Ecology and Evolution. John Wiley and Sons Ltd; 2020 [cited 2020 Jul 27]. p. 6144–62. Available from: https://onlinelibrary.wiley.com/doi/full/10.1002/ece3.6301

6. Nicol S. Krill, Currents, and Sea Ice: Euphausia superba and Its Changing Environment [Internet]. Vol. 56, BioScience. Oxford Academic; 2006 Feb [cited 2020 Aug 3]. Available from: www.biosciencemag.org

7. Hofmann EE, Murphy EJ. Advection, krill, and Antarctic marine ecosystems. Antarct Sci. 2004;16(4):487–99.

8. Stammerjohn S, Massom R, Rind D, Martinson D. Regions of rapid sea ice change: An inter-hemispheric seasonal comparison. Geophys Res Lett. 2012;39(6):1–8.

9. Kawaguchi S, Ishida A, King R, Raymond B, Waller N, Constable A, et al. Risk maps for Antarctic krill under projected Southern Ocean acidification. Nat Clim Chang [Internet]. 2013 [cited 2020 Aug 3];3(9):843–7. Available from: www.nature.com/natureclimatechange

10. Meredith MP, King JC. Rapid climate change in the ocean west of the Antarctic Peninsula during the second half of the 20th century. Geophys Res Lett. 2005;32(19): 1–5.

11. Teschke M, Wendt S, Kawaguchi S, Kramer A, Meyer B. A circadian clock in antarctic krill: An endogenous timing system governs metabolic output rhythms in the euphausid species Euphausia superba. PLoS One. 2011;6(10).

12. Meyer B, Teschke M. Physiology of Euphausia superba [Internet]. Volume 1. Siegel V, editor. Kiel, Germany: Springer; 2016. 145–174 p. Available from: http://www.springer.com/series/10290

13. Reilly S, Hedley S, Borberg J, Hewitt R, Thiele D, Watkins J, et al. Biomass and energy transfer to baleen whales in the South Atlantic sector of the Southern Ocean. Deep Sea Res Part II Top Stud Oceanogr. 2004;51(12–13):1397–409.

14. Ware C, Wiley DN, Friedlaender AS, Weinrich M, Hazen EL, Bocconcelli A, et al. Bottom side-roll feeding by humpback whales (*Megaptera novaeangliae*) in the southern Gulf of Maine, U.S.A. Mar Mammal Sci. 2014;30(2):494–511.

15. Watt CA, Ferguson SH. Fatty acids and stable isotopes (δ^13^ C and δ^15^ N) reveal temporal changes in narwhal (*Monodon monoceros*) diet linked to migration patterns. Mar Mammal Sci [Internet]. 2015 Jan 1 [cited 2020 Aug 28];31(1):21–44. Available from: http://doi.wiley.com/10.1111/mms.12131

16. Bridge ES, Kelly JF, Xiao X, Takekawa JY, Hill NJ, Yamage M, et al. Bird migration and avian influenza: A comparison of hydrogen stable isotopes and satellite tracking methods. Ecol Indic. 2014 Oct 1;45:266–73.

17. Robillard A, Gauthier G, Therrien J-F, Fitzgerald G, Provencher JF, Bêty J. Variability in stable isotopes of snowy owl feathers and contribution of marine resources to their winter diet. J Avian Biol [Internet]. 2017 Jun 1 [cited 2020 Aug 28];48(6):759–69. Available from: http://doi.wiley.com/10.1111/jav.01257

18. Kaczensky P, Šturm MB, Sablin M V, Voigt CC, Smith S, Ganbaatar O, et al. Stable isotopes reveal diet shift from pre-extinction to reintroduced Przewalski’s horses. Nature [Internet]. 2017 [cited 2020 Aug 28];7(5950). Available from: www.nature.com/scientificreports/

19. Chiaradia A, Ramírez F, Forero MG, Hobson KA. Stable Isotopes (δ13C, δ15N) Combined with Conventional Dietary Approaches Reveal Plasticity in Central-Place Foraging Behavior of Little Penguins Eudyptula minor. Front Ecol Evol [Internet]. 2016 Jan 11 [cited 2020 Aug 28];3(JAN):154. Available from: http://journal.frontiersin.org/Article/10.3389/fevo.2015.00154/abstract

20. Borrell A, Gómez-Campos E, Aguilar A. Influence of Reproduction on Stable-Isotope Ratios: Nitrogen and Carbon Isotope Discrimination between Mothers, Fetuses, and Milk in the Fin Whale, a Capital Breeder. Physiol Biochem Zool [Internet]. 2016 Jan 1 [cited 2020 Aug 28];89(1):41–50. Available from: https://www.journals.uchicago.edu/doi/10.1086/684632

21. Pethybridge H, Choy CA, Logan JM, Allain V, Lorrain A, Bodin N, et al. A global meta-analysis of marine predator nitrogen stable isotopes: Relationships between trophic structure and environmental conditions. Glob Ecol Biogeogr [Internet]. 2018 Sep 1 [cited 2020 Aug 28];27(9):1043–55. Available from: http://doi.wiley.com/10.1111/geb.12763

22. Connolly RM, Waltham NJ. Spatial analysis of carbon isotopes reveals seagrass contribution to fishery food web. Ecosphere [Internet]. 2015 Sep 1 [cited 2020 Aug 28];6(9):art148. Available from: http://doi.wiley.com/10.1890/ES14-00243.1

23. Eisenmann P, Fry B, Holyoake C, Coughran D, Nicol S, Bengtson Nash S. Isotopic evidence of a wide spectrum of feeding strategies in Southern hemisphere humpback whale baleen records. PLoS One. 2016;11(5):1–20.

24. Schwarz D, Spitzer SM, Thomas AC, Kohnert CM, Keates TR, Acevedo-Gutiérrez A. Large-scale molecular diet analysis in a generalist marine mammal reveals male preference for prey of conservation concern. Ecol Evol [Internet]. 2018 Oct 1 [cited 2020 Aug 28];8(19):9889–905. Available from: https://onlinelibrary.wiley.com/doi/full/10.1002/ece3.4474

25. Acevedo J, Haro D, Dalla Rosa L, Aguayo-Lobo A, Hucke-Gaete R, Secchi E, et al. Evidence of spatial structuring of eastern South Pacific humpback whale feeding grounds. Endanger Species Res. 2013;22(1):33–8.

26. Witteveen BH, Worthy GAJ, Foy RJ, Wynne KM. Modeling the diet of humpback whales: An approach using stable carbon and nitrogen isotopes in a Bayesian mixing model. Mar Mammal Sci. 2012;28(3):233–50.

27. Zuev AG, Rozanova OL, Tsurikov SM, Panchenko PL, Ershova MA, Smolyarova DD, et al. Stable Isotope Trophic Fractionation (^13^C/^12^C and ^15^N/^14^N) in Mycophagous Diptera Larvae. Biol Bull. 2019;46(5):457–65.

28. Mill AC, Pinnegar JK, Polunin NVC. Explaining isotope trophic-step fractionation: Why herbivorous fish are different. Funct Ecol. 2007;21(6):1137–45.

29. Fry B. Stable isotope ecology. New York: Springer Science + Buisness Media, LLC; 2008. 1–16 p.

30. DeNiro MJ, Epstein S. Influence of diet on the distribution of carbon isotopes in animals. Microw Opt Technol Lett. 1978;42:495–506.

31. Minagawa M, Wada E. Stepwise enrichment of ^15^N along food chains: Further evidence and the relation between δ^15^N and animal age. Geochim Cosmochim Acta. 1984;48(5): 1135–40.

32. Trites AW. Marine mammal trophic levels and trophic interactions [Internet]. 3rd ed. Encyclopedia of Ocean Sciences. Elsevier Ltd.; 2019. 589–594 p. Available from: http://dx.doi.org/10.1016/B978-0-12-409548-9.11618-5

33. Seyboth E, Botta S, Mendes CRB, Negrete J, Dalla Rosa L, Secchi ER. Isotopic evidence of the effect of warming on the northern Antarctic Peninsula ecosystem. Deep Res Part II Top Stud Oceanogr [Internet]. 2018;149(December 2017):218–28. Available from: https://doi.org/10.1016/j.dsr2.2017.12.020

34. Altabet MA, Francois R. Sedimentary nitrogen isotopic ratio as a recorder for surface ocean nitrate utilization. Global Biogeochem Cycles. 1994;8(1):103–16.

35. Goericke R, Fry B. Variations of marine plankton in δ^13^N with latitude, temperature, and dissolved CO2 in the world ocean. Glob Biochem Cycles. 1994;8(1):85–90.

36. Wada E, Terazaki M, Kabaya Y, Nemoto T. ^15^N and ^13^C abundances in the Antartic Ocean with emphasis on the biogeochemical structure of the food web. Deep Sea Res Part A, Oceanogr Res Pap. 1987;34(5–6):829–41.

37. Cherel Y. Isotopic niches of emperor and Adélie penguins in Adélie Land, Antarctica. 2008;813–21.

38. Hall-Aspland SA, Hall AP, Rogers TL. A new approach to the solution of the linear mixing model for a single isotope: Application to the case of an opportunistic predator. Oecologia. 2005;143(1):143–7.

39. Hodum PJ, Hobson KA. Trophic relationships among Antarctic fulmarine petrels : insights into dietary overlap and chick provisioning strategies inferred from stable-isotope (δ^15^N and δ^13^C) analysis. Mar Ecol Prog Ser. 2000;198:273–81.

40. Davenport SR, Bax NJ. A trophic study of a marine ecosystem off southeastern Australia using stable isotopes of carbon and nitrogen. Can J Fish Aquat Sci. 2002;59(3):514–30.

41. Harris BP, Young JW, Revill AT, Taylor MD. Understanding diel-vertical feeding migrations in zooplankton using bulk carbon and nitrogen stable isotopes. J Plankton Res. 2014;36(4):1159–63.

42. Hobson KA, Welch HE. Cannibalism and trophic structure in a high Arctic lake: insights from stable-isotope analysis. Can J Fish Aquat Sci. 1995;52(6): 1195–201.

43. Ponsard S, Averbuch P. Should growing and adult animals fed on the same diet show different δ^15^N values? Rapid Commun Mass Spectrom. 1999;13(13):1305–10.

44. Overman NC, Parrish DL. Stable isotope composition of walleye: ^15^N accumulation with age and area-specific differences in δ^13^C. Can J Fish Aquat Sci. 2001;58(6): 1253–60.

45. Vanderklift MA, Ponsard S. Sources of variation in consumer-diet δ^15^N enrichment: A meta-analysis. Oecologia. 2003;136(2):169–82.

46. Cherry SG, Derocher AE, Hobson KA, Stirling I, Thiemann GW. Quantifying dietary pathways of proteins and lipids to tissues of a marine predator. J Appl Ecol. 2011;48(2):373–81.

47. Tieszen LL, Boutton TW, Tesdahl KG, Slade NA. Fractionation and turnover of stable carbon isotopes in animal tissues: Implications for δ^13^C analysis of diet. Oecologia. 1983;57(1–2):32–7.

48. Budge SM, Iverson SJ, Koopman HN. Studying trophic ecology in marine ecosystems using fatty acids: A primer on analysis and interpretation. Mar Mammal Sci. 2006;22(4):759–801.

49. Noren DP, Mocklin JA. Review of cetacean biopsy techniques: Factors contributing to successful sample collection and physiological and behavioral impacts. Mar Mammal Sci. 2012;28(1):154–99.

50. Castrillon J, Huston W, Bengtson Nash S. The blubber adipocyte index: A nondestructive biomarker of adiposity in humpback whales (*Megaptera novaeangliae*). Ecol Evol. 2017;7(14):5131–9.

51. Filatova OA, Witteveen BH, Goncharov AA, Tiunov A V., Goncharova MI, Burdin AM, et al. The diets of humpback whales (*Megaptera novaeangliae*) on the shelf and oceanic feeding grounds in the western North Pacific inferred from stable isotope analysis. Mar Mammal Sci. 2013;29(3):253–65.

52. Todd S, Ostrom P, Lien J, Abrajano J. Use of biopsy samples of humpback whale (*Megaptera novaeangliae*) skin for stable isotope (δ^13^C) determination. J Northwest Atl Fish Sci. 1997;22(December 1997):71–6.

53. Rolff C, Elmgren R. Use of riverine organic matter in plankton food webs of the Baltic Sea. Mar Ecol Prog Ser. 2000;197:81–101.

54. Post DM, Layman CA, Arrington DA, Takimoto G, Quattrochi J, Montaña CG. Getting to the fat of the matter: Models, methods and assumptions for dealing with lipids in stable isotope analyses. Oecologia. 2007;152(1):179–89.

55. Ryan C, McHugh B, Trueman CN, Harrod C, Berrow SD, O’Connor I. Accounting for the effects of lipids in stable isotope (δ^13^C and δ^15^N values) analysis of skin and blubber of balaenopterid whales. Rapid Commun Mass Spectrom. 2012;26(23):2745–54.

56. Folch J, Lees M, Sloane Stanley GH. A simple method for the isolation and purification of total lipides from animal tissues. J Biol Chem. 1957;226(1):497–509.

57. Bligh EG, Dyer WJ. A rapid method of total lipid extraction and purification. Can J Biochem Physiol. 1959;37(8).

58. Pethybridge HR, Parrish CC, Bruce BD, Young JW, Nichols PD. Lipid, fatty acid and energy density profiles of white sharks: Insights into the feeding ecology and ecophysiology of a complex top predator. PLoS One. 2014;9(5).

59. Groß J, Fry B, Burford M., Bengtson Nash S. Accounting for lipid interference when evaluating diet via stable isotope values in skin and blubber of southern hemisphere humpback whales. (1).

60. Fry B. Stable isotopic indicators of habitat use by Mississippi River Fish. J North Am Benthol Soc. 2002;21(4):676–85.

61. R Core Team. Integrated development for R. [Internet]. Vol. 42, RStudio. Boston, MA: RStudio PBC; 2020. p. 14. Available from: https://rstudio.com

62. GraphPad S. GraphPad Prism [Internet]. San Diego, California: GraphPad; 2020. Available from: www.graphpad.com

63. Witteveen BH, Worthy GAJ, Wynne KM, Hirons AC, Andrews AG, Markel RW. Trophic levels of North Pacific Humpback whales (*Megaptera novaeangliae*) through analysis of stable isotopes: Implications on prey and resource quality. Aquat Mamm. 2011;37(2): 101–10.

64. Haro D, Sabat P, Arreguín-Sánchez F, Neira S, Hernández-Padilla J. Trophic role of the humpback whale (*Megaptera novaeangliae*) in the feeding area of Magellan Strait, Chile. Ecol Indic. 2020;109:105796.

65. Paterson RA, Paterson P, Cato DH. Status of humpback whales, *Megaptera novaeangliae*, in east Australia at the end of the 20th century. Mem MUSEUM. 2001;2(47):579–86.

66. Chittleborough RG. Dynamics of two populations of the humpback whale. *Megaptera novaeangliae* (borowski). Mar Freshw Res. 1965;16(1):33–128.

67. Polito MJ, Reiss CS, Trivelpiece WZ, Patterson WP, Emslie SD. Stable isotopes identify an ontogenetic niche expansion in Antarctic krill (*Euphausia superba*) from the South Shetland Islands, Antarctica. Mar Biol. 2013;160(6):1311–23.

68. Chikaraishi Y, Ogawa NO, Kashiyama Y, Takano Y, Suga H, Tomitani A, et al. Determination of aquatic food-web structure based on compound-specific nitrogen isotopic composition of amino acids. Limnol Oceanogr Methods. 2009;7(NOV):740–50.

69. Chikaraishi Y, Kashiyama Y, Ogawa NO, Kitazato H, Ohkouchi N. Metabolic control of nitrogen isotope composition of amino acids in macroalgae and gastropods: Implications for aquatic food web studies. Mar Ecol Prog Ser. 2007;342:85–90.

70. McClelland JW, Montoya JP. Trophic relationships and the nitrogen isotopic composition of amino acids in plankton. Ecology. 2002;83(8):2173–80.

71. Zilversmith D., Entenman C, Fishler C. On the calculation of “turnover time” and “turnover rate” from experiments involving the use of labeling agents. Gen Physiol. 1942;325–31.

72. Reiner JM. The study of metabolic turnover rates by means of isotopic tracers: I. Fundamental relations. Arch Biochem Biophys. 1953;46(1):53–79.

73. Newsome SD, Clementz MT, Koch PL. Using stable isotope biogeochemistry to study marine mammal ecology. Mar Mammal Sci. 2010;26(3):509–72.

74. Browning NE, Dold C, I-Fan J, Worthy GAJ. Isotope turnover rates and diet-tissue discrimination in skin of ex situ bottlenose dolphins (*Tursiops truncatus*). J Exp Biol. 2014;217(2):214–21.

75. Busquets-Vass G, Newsome SD, Calambokidis J, Serra-Valente G, Jacobsen JK, Aguíñiga-García S, et al. Estimating blue whale skin isotopic incorporation rates and baleen growth rates: Implications for assessing diet and movement patterns in mysticetes. PLoS One [Internet]. 2017 [cited 2020 Aug 28];5(12). Available from: https://doi.org/10.1371/journal.pone.0177880

76. Wolf N, Newsome SD, Peters J, Fogel ML. Variability in the routing of dietary proteins and lipids to consumer tissues influences tissue-specific isotopic discrimination. Rapid Commun Mass Spectrom. 2015;29(15):1448–56.

77. Caut S, Angulo E, Courchamp F. Variation in discrimination factors (Δ^15^N and Δ^13^C): The effect of diet isotopic values and applications for diet reconstruction. J Appl Ecol. 2009;46(2):443–53.

78. Misra BB, Mariel RHI, Ivonne HBG, Emanuel HN, Raúl DG, Cristina CDR. 1H NMR metabolomic analysis of skin and blubber of bottlenose dolphins reveal a functional metabolic dichotomy. Comp Biochem Physiol - Part D Genomics Proteomics [Internet]. 2019;30(February):25–32. Available from: https://doi.org/10.1016/j.cbd.2019.02.004

79. McConnaughey T, McRoy CP. Food-Web structure and the fractionation of Carbon isotopes in the bering sea. Mar Biol. 1979;53(3):257–62.

80. Logan JM, Lutcavage ME. A comparison of carbon and nitrogen stable isotope ratios of fish tissues following lipid extractions with non-polar and traditional chloroform/methanol solvent systems. Rapid Commun Mass Spectrom. 2008;22:1081–6.

81. Sotiropoulos MA, Tonn WM, Wassenaar LI. Effects of lipid extraction on stable carbon and nitrogen isotope analyses of fish tissues: Potential consequences for food web studies. Ecol Freshw Fish. 2004;13(3):155–60.

82. Yurkowski DJ, Hussey NE, Semeniuk C, Ferguson SH, Fisk AT. Effects of lipid extraction and the utility of lipid normalization models on δ^13^C and δ^15^N values in Arctic marine mammal tissues. Polar Biol. 2015;38(2):131–43.

83. Bearhop S, Waldron S, Furness RW. Influence of Lipid and Uric Acid on δ^13^C and δ^15^N Values of Avian Blood: Implications for Trophic Studies. Auk. 2000;117(2):504–7.

84. Logan JM, Jardine TD, Miller TJ, Bunn SE, Cunjak RA, Lutcavage ME. Lipid corrections in carbon and nitrogen stable isotope analyses: Comparison of chemical extraction and modelling methods. J Anim Ecol. 2008;77(4):838–46.

